# A multi-omics digital research object for the genetics of sleep regulation

**DOI:** 10.1101/586206

**Authors:** Maxime Jan, Nastassia Gobet, Shanaz Diessler, Paul Franken, Ioannis Xenarios

## Abstract

More and more researchers make use of multi-omics approaches to tackle complex cellular and organismal systems. It has become apparent that the potential for re-use and integrate data generated by different labs can enhance knowledge. However, a meaningful and efficient re-use of data generated by others is difficult to achieve without in depth understanding of how these datasets were assembled. We therefore designed and describe in detail a digital research object embedding data, documentation and analytics on mouse sleep regulation. The aim of this study was to bring together electrophysiological recordings, sleep-wake behavior, metabolomics, genetics, and gene regulatory data in a systems genetics model to investigate sleep regulation in the BXD panel of recombinant inbred lines. We here showcase both the advantages and limitations of providing such multi-modal data and analytics. The reproducibility of the results was tested by a bioinformatician not implicated in the original project and the robustness of results was assessed by re-annotating genetic and transcriptome data from the mm9 to the mm10 mouse genome assembly.

## Background & Summary

A good night’s sleep is essential for optimal performance, wellbeing and health. Chronically disturbed or curtailed sleep can have long-lasting adverse effects on health with associated increased risk for obesity and type-2 diabetes^1^.

To gain insight into the molecular signaling pathways regulating undisturbed sleep and the response to sleep restriction in the mouse, we performed a population-based multi-level screening known as *systems genetics* ^2^. This approach allows to chart the molecular pathways connecting genetic variants to complex traits through the integration of multiple *omics datasets such as transcriptomics, proteomics, metabolomics or microbiomes ^3^. We built a systems genetics resource based on the BXD panel, a population of recombinant inbred lines of mice ^4^, that has been used for a number of complex traits and *omics screening such as brain slow-waves during NREM sleep ^5^, glucose regulation ^6^, cognitive aging ^7^ and mitochondria proteomics ^8^.

We phenotyped 34 BXD/RwwJ inbred lines, 4 BXD/TyJ, 2 parental strains C57BL6/J and DBA/2J and their reciprocal F1 offspring. Mice of these 42 lines were challenged with 6h of sleep deprivation (SD) to evaluate the effects of insufficient sleep on sleep-wake behavior and brain activity (electroencephalogram or EEG; Figure 1, Experiment 1) and, on gene expression and metabolites (Figure 1, Experiment 2). For Experiment 1 we recorded the EEG together with muscle tone (electromyogram or EMG) and locomotor activity (LMA) continuously for 4 days. Based on the EEG/EMG signals we determined sleep-wake state [wakefulness, rapid-eye movement (REM) sleep, and non-REM (NREM) sleep] as well as the spectral composition of the EEG signal as end phenotypes. For Experiment 2 we quantified mRNA levels in cerebral cortex and liver using illumina HiSeq 2500 RNA-sequencing and performed a targeted metabolomics screen on blood using Biocrates p180 liquid chromatography (LC-) and Flow injection analysis (FIA-) coupled with mass spectrometry (MS). These transcriptome and metabolome data are regarded as intermediate phenotypes linking genome information to the sleep-wake related end phenotypes.

**Figure 1:**
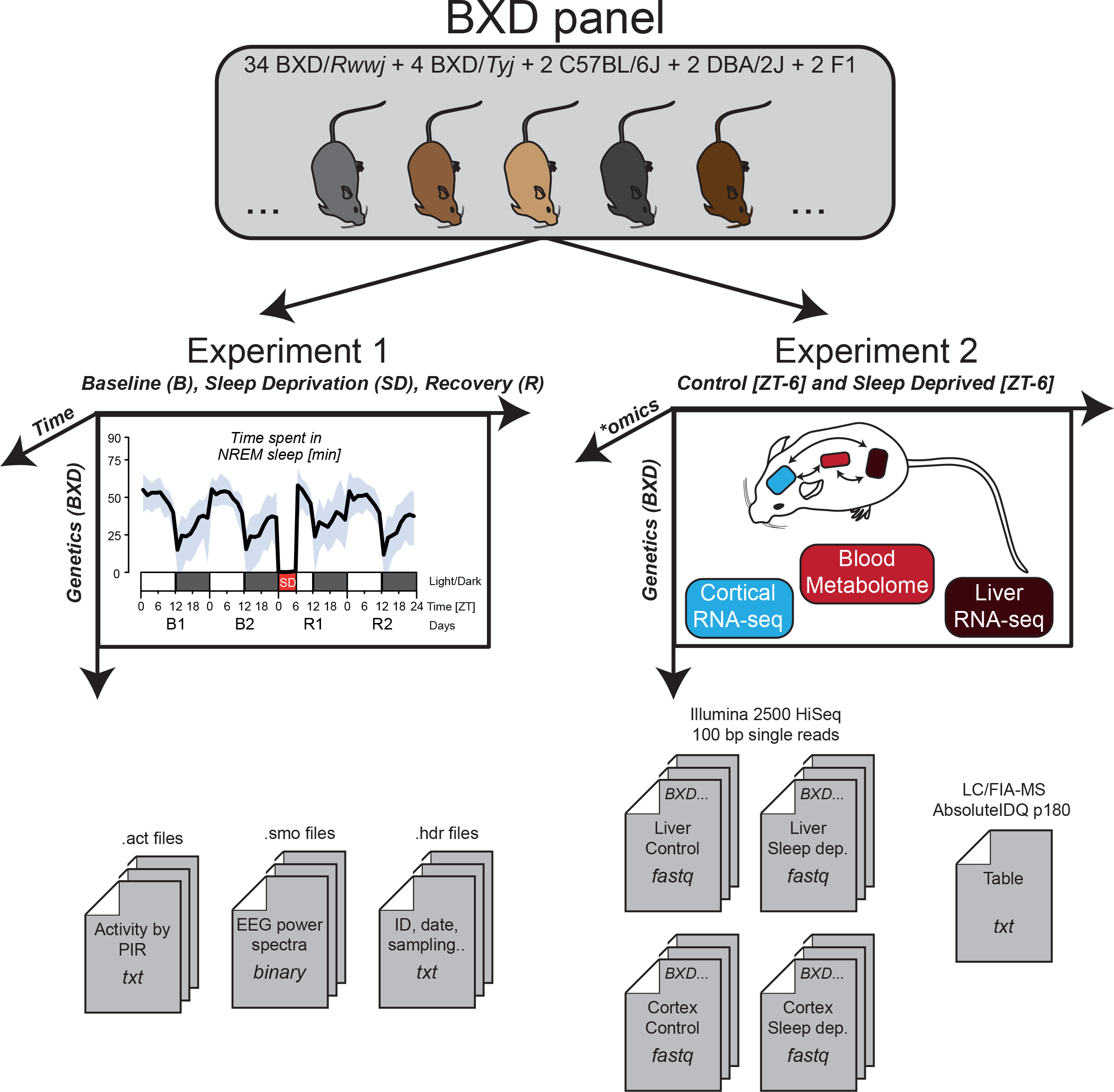
Data generation. The behavioral/EEG end-phenotypes of the BXD mouse panel were quantified in experiment 1. Mice were recorded for 4 days: 2 days of baseline (B1 & B2), followed by 6h of sleep deprivation (SD) and 2 days of recovery (R1 & R2). EEG spectral composition was written in *.smo* files, activity in *.act* files and meta-data in *.hdr* files. Blood metabolomics, liver transcriptomics and cortical transcriptomics were quantified in experiment 2. ‘Control’ and ‘Sleep deprived’ batches were sampled at a single time point: ZT6 (i.e. directly after sleep deprivation for the ‘sleep deprived’ batch). Transcriptomics was performed on pooled sampled per BXD strains. For blood metabolomics, metabolite quantification was performed for each BXD replicates.

The keystone of systems genetics is data integration. Accordingly, the scientific community can benefit from facilited dataset sharing to integrate the results of their own experiment with that of others. However, reliable methods for data integration are needed and require a broad range of expertise such as in mathematical and statistical models ^9^, computational methods ^10^, visualization strategies ^11^, and deep understanding of complex phenotypes. Therefore, data sharing should not be limited to the dataset *per se* but also to analytics in the form of analysis workflows, code, interpretation of results, and meta-data ^12^. The concept of a Digital Research Object (DRO) was proposed to group dataset and analytics into one united package ^13^. Various guidelines have been suggested to address the challenges of sharing such DRO with the goal to improve and promote the human and computer knowledge sharing, like the FAIR (Findable, Accessible, Interoperable, Reusable) principles proposed by FORCE 11 ^14^ or by the DB2K (Big Data to Knowledge) framework. These guidelines concern biomedical workflow, meta-data structures and computer infrastructures facilitating the reusability and interoperability of digital resources ^15^. Although such guidelines are often described and applied in the context of single data-type assays, they can be challenging to achieve for trans-disciplinary research projects such as systems genetics, in which multiple data types, computer programs, references and novel methodologies need to be combined ^16^. Moreover, applying these principles can also be discouraging because of the time required for new working routines to become fully reproducible ^17^ and because only few biomedical journals have standardized and explicit data-sharing ^18^ or reproducibility ^19^ policies. Nonetheless, DROs are essential for scientific reliability ^20^, and can save time if a dataset or methods specific to a study need to be reused or improved by different users such as colleagues at other institutes, new comers to the lab, or at long-term yourself.

We here complement our previous publication ^2^ by improving the raw and processed data availability. We describe in more details the different bioinformatics steps that were applied to analyze this resource and improve the analytical pipeline reproducibility by generating *R* reports and provide code. Finally, we assess the reproducibility of our bioinformatic pipeline from the perspective of a new student in bioinformatics that recently joined the group, and the robustness of the results by changing both the mouse reference genome and the RNA-seq reads alignment to new standards.

## Methods

These methods are an expanded version of the methods described in our related paper ^2^. Appreciable portions are reproduced verbatim to deliver a complete description of the data and analytics with the aim to enhance reproducibility.

Experiment 1 and Experiment 2 (Figure 1) were approved by the veterinary authorities of the state of Vaud, Switzerland (SCAV authorization #2534).

### Animal, breeding, and housing conditions

34 BXD lines originating from the University of Tennessee Health Science Center (Memphis, TN, United States of America) were selected for Experiment 1 and Experiment 2. These lines were randomly chosen from the newly generated advanced recombinant inbred line (ARIL) RwwJ panel ^4^, although lines with documented poor breeding performance were not considered. 4 additional BXD RI strains were chosen from the older *TyJ* panel for reproducibility purposes and were obtained directly from the Jackson Laboratory (JAX, Bar Harbor, Maine). The names used for some of the BXD lines have been modified over time to reflect genetic proximity. Table 1 lists the BXD line names we used in our files alongside the corresponding current JAX names and IDs. In our analyses, we discarded the BXD63/RwwJ line for quality reasons (see Technical Validation) as well as the 4 older BXD strains that were derived from a different DBA/2 sub-strains, i.e. DBA/2Rj instead of DBA/2J for RwwJ lines ^21^. The methods below describe the remaining 33 BXD lines, F1 and parental strains. Two breeding trios per BXD strain were purchased from a local facility (EPFL-SV, Lausanne, Switzerland) and bred in-house until sufficient offspring was obtained. The parental strains DBA/2J (D2), C57BL6/J (B6) and their reciprocal F1 offspring (B6D2F1 [BD-F1] and D2B6F1 [DB-F1]) were bred and phenotyped alongside. Suitable (age and sex) offspring was transferred to our sleep-recording facility, where they were singly housed, with food and water available ad libitum, at a constant temperature of 25°C and under a 12 h light/12 h dark cycle (LD12:12, fluorescent lights, intensity 6.6 cds/m^2^, with ZT0 and ZT12 designating light and dark onset, respectively). Male mice aged 11–14 week at the time of experiment were used for phenotyping, with a mean of 12 animals per BXD line among all experiments. Note that 3 BXD lines had a lower replicate number (n), with respectively BXD79 (n = 6), BXD85 (n = 5), and BXD101 (n = 4) because of poor breeding success. For the remaining 30 BXD lines, replicates were distributed as follows: for EEG/behavioral phenotyping (Experiment 1 in Fig 1; mean = 6.2/line; 5 ≤ n ≤ 7) and for molecular phenotyping (Experiment 2 in Fig 1; mean = 6.8/line; 6 ≤ n ≤ 9). Additionally, to control for the reproducibility of the outcome variables over the experiment, parental lines were phenotyped twice—i.e., at the start (labeled in files as B61 and DB1) and end (labeled B62 and DB2) of the breeding and data-collecting phase, which spanned 2 years (March 2012–December 2013). To summarize, distributed over 32 experimental cohorts, 227 individual mice were used for behavioral/EEG phenotyping (Experiment 1) and 263 mice for tissue collection for transcriptome and metabolome analyses (Experiment 2), the latter being divided into sleep deprived (SD) and controls (“Ctr”; see Study design section below). We randomized the lines across the experimental cohorts so that biological replicates of 1 line were collected/recorded on more than 1 occasion while also ensuring that an even number of mice per line was included for tissue collection so as to pair SD and “Ctr” individuals within each cohort (for behavioral/EEG phenotyping, each mouse serves as its own control).

**Table 1:** Names of BXD lines used in our files with the corresponding JAX name and ID. F1 lines (DB and BD) were generated in house. BXD line names in our files can also be found without ‘0’ i.e. BXD50 instead of BXD050. Further note that the names we used followed an older nomenclature and some names therefore differ from the current JAX names listed.

| Name in files | JAX Name | JAX ID |
| --- | --- | --- |
| BXD005 | BXD5/TyJ | 000037 |
| BXD029TL / BXD029t | BXD29- Tlr4<sup>lps-2J</sup>/J | 000029 |
| BXD029 | BXD29/Ty | 010981 |
| BXD032 | BXD32/TyJ | 000078 |
| BXD043 | BXD43/RwwJ | 007093 |
| BXD044 | BXD44/RwwJ | 007094 |
| BXD045 | BXD45/RwwJ | 007096 |
| BXD048 | BXD48/RwwJ | 007097 |
| BXD049 | BXD49/RwwJ | 007098 |
| BXD050 | BXD50/RwwJ | 007099 |
| BXD051 | BXD51/RwwJ | 007100 |
| BXD055 | BXD55/RwwJ | 007103 |
| BXD056 | BXD56/RwwJ | 007104 |
| BXD061 | BXD61/RwwJ | 007106 |
| BXD063 | BXD63/RwwJ | 007108 |
| BXD064 | BXD64/RwwJ | 007109 |
| BXD065 | BXD65/RwwJ | 007110 |
| BXD066 | BXD66/RwwJ | 007111 |
| BXD067 | BXD67/RwwJ | 007112 |
| BXD070 | BXD70/RwwJ | 007115 |
| BXD071 | BXD71/RwwJ | 007116 |
| BXD073 | BXD73/RwwJ | 007117 |
| BXD075 | BXD75/RwwJ | 007119 |
| BXD079 | BXD79/RwwJ | 007123 |
| BXD081 | BXD81/RwwJ | 007125 |
| BXD083 | BXD83/RwwJ | 007126 |
| BXD084 | BXD84/RwwJ | 007127 |
| BXD085 | BXD85/RwwJ | 007128 |
| BXD087 | BXD87/RwwJ | 007130 |
| BXD089 | BXD89/RwwJ | 007132 |
| BXD090 | BXD90/RwwJ | 007133 |
| BXD095 | BXD95/RwwJ | 007138 |
| BXD096 | BXD48a/RwwJ | 007139 |
| BXD097 | BXD65a/RwwJ | 007140 |
| BXD098 | BXD98/RwwJ | 007141 |
| BXD100 | BXD100/RwwJ | 007143 |
| BXD101 | BXD101/RwwJ | 007144 |
| BXD103 | BXD73b/RwwJ | 007146 |
| C57BL6 / B61 / B62 | C57BL/6J | 000664 |
| DBA2 / DB1 / DB2 | DBA/2J | 000671 |
| DB / DXB F1 / DBA/2J x<br>C57BL6/J F1 | - | - |
| BD / BXD F1 / C57BL6/J x<br>DBA/2J F1 | - | - |

### Study design

The study consisted of 2 experiments, i.e., Experiments 1 and 2 (Figure 1). Animals of both experiments were maintained under the same housing conditions. Animals in Experiment 1 underwent surgery and, after a >10 days recovery period, electroencephalography (EEG), electromyography (EMG) and locomotor activity (LMA) were recorded continuously for a 4-day period starting at ZT0. The first 2 days were considered Baseline (B1 and B2). The first 6 hours of Day 3 (ZT0–6), animals were sleep deprived (SD) in their home cage by “gentle handling” referring to preventing sleep by changing litter, introducing paper tissue, present a pipet near the animal or gently tapping the cage. Experimenters performing the SD rotated every 1 or 2 hours for the SD duration (for more information, see ^22^). The remaining 18 h of Day 3 and the entire Day 4 were considered Recovery (R1 and R2).

Half of the animals included in Experiment 2 underwent SD alongside the animals of Experiment 1. The other half was left undisturbed in another room (i.e., control or Ctr, also referred as Non Sleep Deprived or NSD). Both SD and “Ctr” mice of Experiment 2 were killed at ZT6 (i.e., immediately after the end of the SD) for sampling of liver and cerebral cortex tissue as well as trunk blood. All mice were left undisturbed for at least 2 days prior to SD.

### Experiment 1: EEG/EMG and LMA recording and signal pre-processing

EEG/EMG surgery was performed under deep anesthesia. IP injection of Xylazine/Ketamine mixture (91/14.5 mg/kg, respectively) ensures a deep plane of anesthesia for the duration of the surgery (i.e., around 30 min). Analgesia was provided the evening prior and the 3 day after surgery with Dafalgan in the drinking water (200–300 mg/kg). Six holes were drilled into the cranium, 4 for screws to fix the connector with Adhesive Resin Cement, 2 for electrodes. The caudal electrode was placed over the hippocampal structure and the rostral electrode was placed over the frontal cerebral cortex. Two gold-wire electrode were inserted into the neck muscle for EMG recording (for details, see ^22^). Mice were allowed to recover for at least 10 days prior to baseline recordings. EEG and EMG signals were amplified, filtered, digitized, and stored using EMBLA (Medcare Flaga, Thornton, CO, USA) hardware (A10 recorder) and software (Somnologica). Digitalization of the signal was performed as followed: the analog to digital conversion of the signal was performed at a rate of 2000 Hz, the signal was down sampled at 200 Hz, high-pass filter at 0.0625 Hz was applied to reject DC signal and a notch filter applied at 50 Hz for interfering signals filtering. Signal was then transformed by Discrete Fourier Transform (DFT) to yield power spectra between 0 and 100 Hz with a 0.25 frequency resolution using a 4-seconds time resolution (called an epoch). EEG frequency bins with artefacts of known (line artefacts between 45-55 Hz) and unknown (75-77 Hz) source were removed from the average EEG spectra of all mice. Other specific 0.25 Hz bins containing artefacts (notably the 8.0, 16.0 and 32.0 Hz bins) of unknown source, were removed from individual mice based on the visual inspection of individual EEG spectra in each of the three sleep-wake states (i.e. wakefulness, REM sleep and NREM sleep). Power density in frequency bins deemed artefacted were estimated by linear interpolation. For details, see Pascal scripts in (Data Citation 4, *gitlab* <u>Systems_Genetics_of_Sleep_Regulation</u>).

LMA was recorded by passive infrared (PIR) sensors (Visonic, Tel Aviv, Israel) at 1 min resolution for the duration of the 4-day experiment, using ClockLab (ActiMetrics, IL, USA). Activity data were made available as *.act* files at Figshare (Data Citation 1: *Figshare* LinkToCome).

Offline, the sleep-wake states wakefulness, REM sleep, and NREM sleep were annotated on consecutive 4-second epochs, based on the EEG and EMG pattern. (see Sleep-wake state annotation section). EEG/EMG power spectra and sleep-wake states annotation were made available as *.smo* files at (Data Citation 1: *Figshare* LinkToCome).

### Experiment 2: Tissue collection and preparation

Mice were killed by decapitation after being anesthetized with isoflurane, and blood, cerebral cortex, and liver were collected immediately. The whole procedure took no more than 5 min per mouse. Blood was collected at the decapitation site into tubes containing 10 ml heparin (2 U/μl) and centrifuged at 4,000 rpm during 5 min at 4°C. Plasma was collected by pipetting, flash-frozen in liquid nitrogen, and stored at −80°C until further use. Cortex and liver were flash-frozen in liquid nitrogen immediately after dissection and were stored at −140°C until further use.

For RNA extraction, frozen samples were homogenized for 45 seconds in 1 ml of QIAzol Lysis Reagent (Qiagen; Hilden, Germany) in a gentleMACS M tube using the gentleMACS Dissociator (Miltenyi Biotec; Bergisch Gladbach, Germany). Homogenates were stored at −80°C until RNA extraction. Total RNA was isolated and purified from cortex using the automated nucleic acid extraction system QIAcube (Qiagen; Hilden, Germany) with the RNeasy Plus Universal Tissue mini kit (Qiagen; Hilden, Germany) and were treated with DNAse. Total RNA from liver was isolated and purified manually using the Qiagen RNeasy Plus mini kit (Qiagen; Hilden, Germany), which includes a step for effective elimination of genomic DNA. RNA quantity, quality, and integrity were assessed utilizing the NanoDrop ND-1000 spectrophotometer (Thermo scientific; Waltham, Massachusetts, USA) and the Fragment Analyzer (Advanced Analytical). The 263 mice initially killed for tissue collection yielded 222 cortex and 222 liver samples of good quality.

Equal amounts of RNA from biological replicates (3 samples per strain, tissue, and experimental condition, except for BXD79, BXD85, and BXD101; see above under Animals, breeding, and housing conditions) were pooled, yielding 156 samples for library preparation. RNA-seq libraries were prepared from 500 ng of pooled RNA using the Illumina TruSeq Stranded mRNA reagents (Illumina; San Diego, California, USA) on a Caliper Sciclone liquid handling robot (PerkinElmer; Waltham, Massachusetts, USA). Libraries were sequenced on the Illumina HiSeq 2500 using HiSeq SBS Kit v3 reagents, with cluster generation using the Illumina HiSeq PE Cluster Kit v3 reagents. Fastq files were pre-processed using the Illumina Casava 1.82 pipeline and bad quality reads tagged with ‘Y’. A mean of 41 M 100 bp single-end reads were obtained (29 M ≤ n ≤ 63 M). Quality of sequences were evaluated using FastQC software (version 0.10.1) and reports made available here (Data Citation 3, bxd.vital-it.ch https://bxd.vital-it.ch/#/dataset/). Figure 2 (A, B, C and D) shows the Phred quality score distribution per base among all samples reads for ‘Cortex Control’, ‘Cortex SD’, ‘Liver Control’ and ‘Liver SD’ respectively. Fastq files were made available at NCBI Gene Expression Omnibus (Data Citation 2: *NCBI Gene Expression Omnibus* GSE112352). Targeted metabolomics analysis was performed using flow injection analysis (FIA) and liquid chromatography/mass spectrometry (LC/MS) as described in ^23,24^. To identify metabolites and measure their concentrations, plasma samples were analyzed using the AbsoluteIDQ p180 targeted metabolomics kit (Biocrates Life Sciences AG, Innsbruck, Austria) and a Waters Xevo TQ-S mass spectrometer coupled to an Acquity UPLC liquid chromatography system (Waters Corporation, Milford, MA, USA). The kit provided absolute concentrations for 188 endogenous compounds from 6 different classes, namely acyl carnitines, amino acids, biogenic amines, hexoses, glycerophospholipids, and sphingolipids. Plasma samples were prepared according to the manufacturer’s instructions. Sample order was randomized, and 3 levels of quality controls (QCs) were run on each 96-well plate. Data were normalized between batches, using the results of quality control level 2 (QC2) repeats across the plate (n = 4) and between plates (n = 4) using Biocrates METIDQ software (QC2 correction). Metabolites below the lower limit of quantification or the limit of detection, as well as above the upper limit of quantification, or with standards out of limits, were discarded from the analysis ^24^. Out of the 188 metabolites assayed, 124 passed these criteria across samples and were used in subsequent analyses. No hexoses were present among the 124 metabolites. Out of the 256 mice killed for tissue collection, 249 plasma samples were used for this analysis. An average of 3.5 animals (3 ≤ n ≤ 6) per line and experimental condition were used (except for BXD79, BXD85, and BXD101 with respectively 2, 1, and 1 animal/condition used; see above under Animals, breeding, and housing conditions). Note that in contrast to the RNA-seq experiment, samples were not pooled but analyzed individually. Mean metabolite levels per BXD lines were made available at bxd.vital-it.ch (Data Citation 3, bxd.vital-it.ch https://bxd.vital-it.ch/#/dataset/), for details see intermediate files (Data Citation 5, *figshare* https://figshare.com/s/51916157a22357755de8).

**Figure 2:**
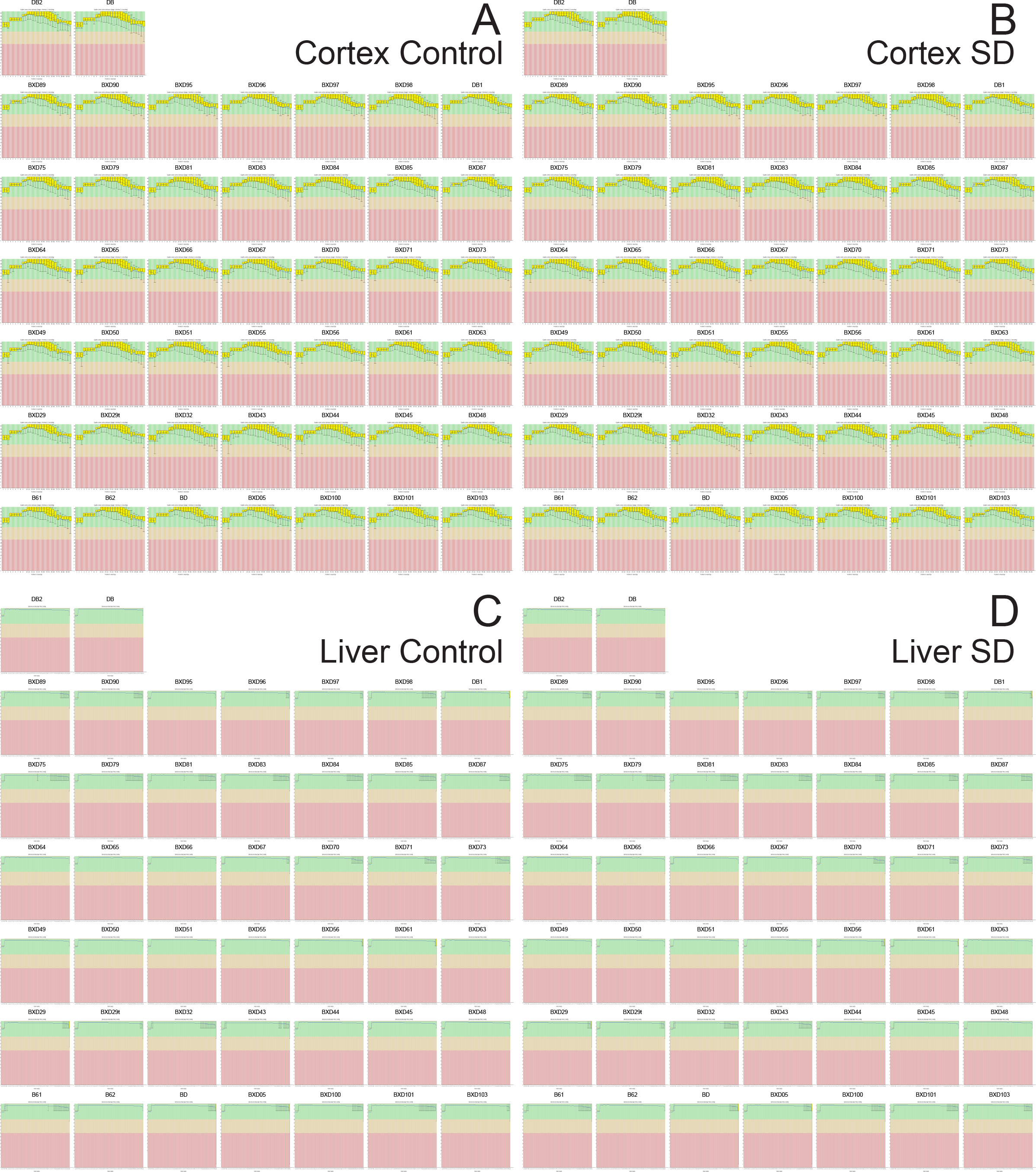
PHRED read quality per base for BXD RNA-sequencing. PHRED quality score based on illumina 1.9. A: Samples from Cortex during control. B: Samples from Cortex after sleep deprivation. C: Samples from Liver during control. D: Samples from Liver after sleep deprivation.

In the same plasma samples, we determined corticosterone levels using an enzyme immunoassay (corticosterone EIA kit; Enzo Life Sciences, Lausanne, Switzerland) according to the manufacturer’s instructions. All samples were diluted 40 times in the provided buffer, kept on ice during the manipulation, and tested in duplicate. BXD lines were spread over multiple 96-well plates in an attempt to control for possible batch effects. In addition, a “control” sample was prepared by pooling plasma from 5 C57BL6/J mice. Aliquots of this control were measured along with each plate to assess plate-to-plate variability. The concentration was calculated in pg/ml based on the average net optical density (at λ = 405 nm) for each standard and sample.

Corticosterone level were made available on figshare (Data Citation 5, *figshare* https://figshare.com/s/51916157a22357755de8)

### Bioinformatics pipeline

To facilitate the interpretation of the complete bioinformatic workflow that was performed on this dataset, we here describe first our general strategy to construct an analytics pipeline with which we hope to improve reproducibility. We then describe the specific methods used to analyze this dataset.

The analytics and input datasets were separated into 3 layers according to increasing level of data abstraction (Figure 3). This hierarchical structure of the workflow was particularly useful to identify steps downstream novel versions of a script or data (e.g. Figure 3, red) and simplify workflow description. The first *low-level* layer contains the procedures needed to reduce and transform the raw-data (i.e. RNA-seq reads, EEG/EMG signals) into an exploitable signal such as sleep phenotypes, genes expression or mice genotypes by further analytical steps. This layer is characterized by long and computationally intensive procedures which required the expertise of different persons, each with their own working environment and preferred informatics language.

**Figure 3:**
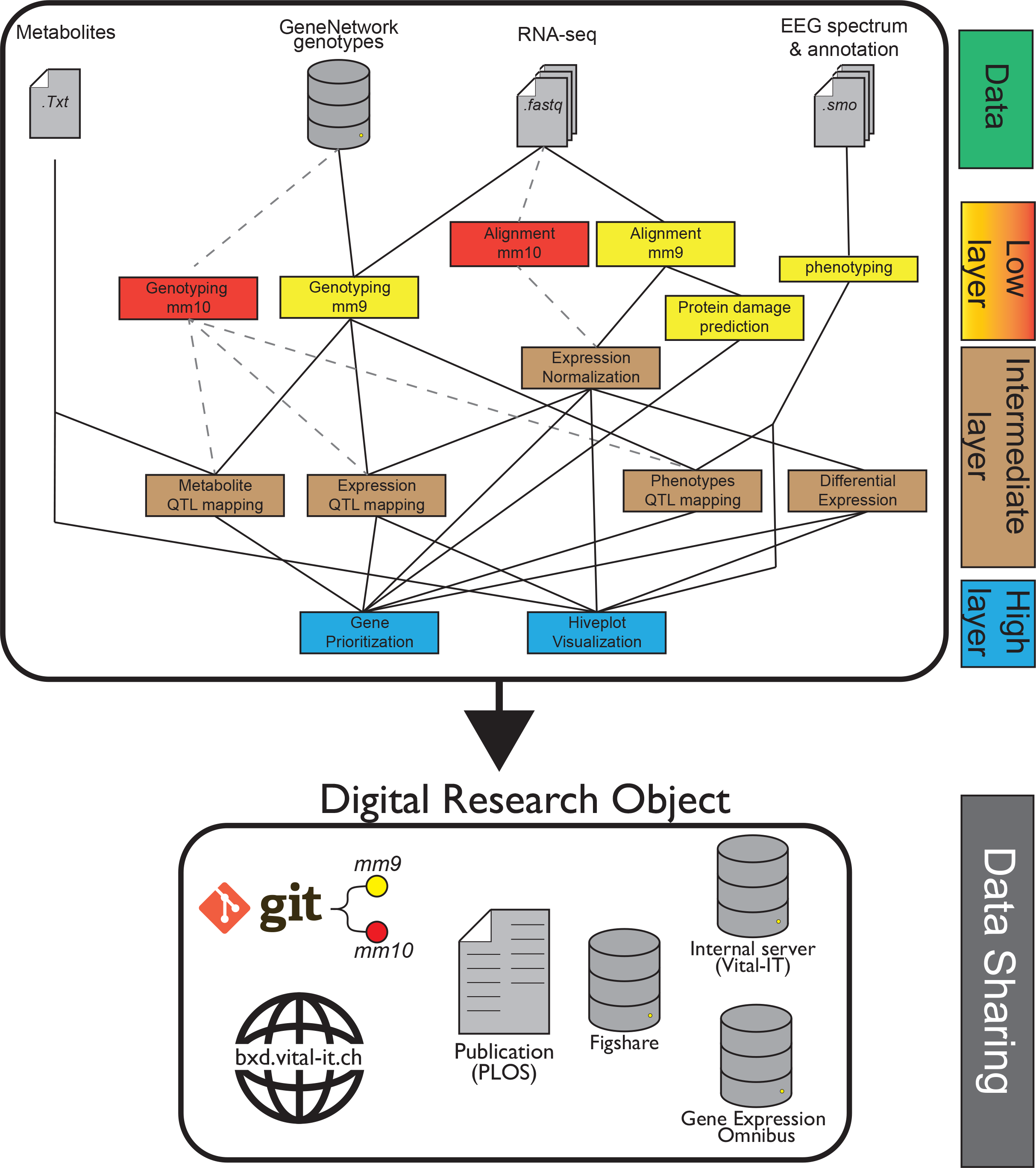
Summary of the bioinformatic analytical pipeline. Representation of the main bioinformatics methods used. Original analyses were performed using the mm9 mouse assembly (yellow). Results were also reproduced using the mm10 mouse assembly (red) and all downstream analyses. Layers represent the scripts organization on gitlab and available intermediate files.

The *intermediate* layer contains some established analyses that could be performed on the data such as gene expression normalization followed by differential expression or Quantitative Trait Locus (QTL) mapping. With the scripts of this layer we explored the effects of sleep deprivation, genetic variations, as well as their interaction on EEG/behavioral phenotypes and intermediate phenotypes.

The *high-level* layer contains the novel integrative methods that we developed to prioritize genes driving sleep regulation and to visually represent the meta-dimensional multi-omics networks underlying sleep phenotypes.

### Code availability on Git

The scripts used for analytics were made available on gitlab (Data Citation 4, *gitlab* Systems_Genetics_of_Sleep_Regulation). The master branch contains the scripts used for our publication and mm9 analysis. A second branch was created for analysis performed on a mm10 mouse references (see Technical Validation). The intermediate files required to run these scripts were made available here: (Data Citation 5, *figshare* https://figshare.com/s/51916157a22357755de8). Finally, a documentation file was generated to understand the hierarchical relationship between the scripts and datasets in a form of a dynamic html document (see Workflow documentation).

### Standard and non-standard semantics

To improve the reproducibility and reusability of our workflow, we tried to prioritize standard semantics and well-established pipelines when it was applicable, such as the RNA-seq processing by STAR and htseq-count ^25^. The use of curated symbols for genes nomenclature by RefSeq allowed a better semantic interoperability with other resources such as Uniprot protein ID using solutions like biomaRt ^26^. We provided some of the references files used in these scripts, like the RefSeq *.gtf* reference file. These annotations can be updated and possibly change the gene quantification with updated version or different genome reference (see References_Files in Data Citation 5, *figshare* https://figshare.com/s/51916157a22357755de8).

However, some steps could not be performed using standards. The EEG/behavioral phenotyping procedure could not be performed by any standard computational workflow or common semantics as none exist. The nomenclature that was chosen in this case to generate unique phenotypic ID was a combination of the phenotype observed (e.g. EEG power during NREM sleep) and the features observed in this phenotype (e.g. delta band 1-4 Hz). These phenotypes were also present as file name and column name in our dataset (Data Citation 5, *figshare* https://figshare.com/s/51916157a22357755de8).

### Favor *R* and *Rmarkdown* reports for reproducible results

Once the data processed within the *low-level* layer, the effect of sleep deprivation, genetics and their interaction were measured using different statistical models and computational methods. We chose to prioritize the programming language *R* as it was the best suited tool for these statistical analyses and for the generation of figures. Beside the advantages of a license-free and portable language, *R* was already recommended as main tool for systems genetics analysis ^27^. Many available packages were particularly adapted for the systems genetics design, involving phenotype-genotype association (*r/qtl*), network analysis (*WGCNA*, *SANTA*, *igraph*), differential expression (*EdgeR*, *DESeq*, *limma*), bayesian network learning (*bnlearn*), visualization (*ggplot2*, *grid*), enrichment (*topGO*, *topAnat*) and parallel computing (*parallel*). Only a few analyses were performed using other softwares, principally for efficiency reasons in cis-/trans-eQTL analysis where the number of models to test was quite large ^28,29^. R is one of the flagships of open science and reproducibility ^30^ with a reviewable source code and the possibility of generating reports known as ‘*Rmarkdown*’ with 2 packages: *knitr* ^31^ and *rmarkdown* ^32^. This report format contains combination of code, figures, and comments within a single *markdown* document that can be easily converted into *pdf* or *html* format. Rmarkdown scripts were made available on (Data Citation 4, *gitlab* Systems_Genetics_of_Sleep_Regulation) and the reports in the form of .html document were made available with data on (Data Citation 5, *figshare* https://figshare.com/s/51916157a22357755de8). To avoid the need to copy/paste some functions shared between *Rmarkdowns* but still display them in our reports, we used the *readLines()* function within Rmarkdown chunks. Finally, the use of the *sessionInfo()* function at the end of the document allowed to keep track of the packages version and the environment variable used. Some of these Rmarkdown reports were generated on a remote cluster instead of the more traditional Rstudio environment, for more information on how to generate these Rmarkdown, see the Usage Notes.

### Workflow documentation

This systems genetics approach was an integrative project that implicated multiple collaborators, that each contributed to the final results, with their own working habit related to their area of expertise. For better reproducibility of the generated files, a critical goal was to keep track of the different files created, associated documents or analytical steps that were produced. For example, EEG/behavioral phenotypes could be found within many files and reports, from *low-level* to *high-level* layers, but their nomenclatures were still hard to interpret as mentioned above, for those not directly related to this project. A newcomer in this project should be able to easily recover the metadata document containing all the physiological phenotypes information (i.e. understand that a metadata document was created and where to find it or who to ask for it) and understand which scripts were used to produce these phenotypes. To establish what was exactly performed, we generated a documentation file containing the essential information and relationships between all the files, scripts, Rmarkdown, small workflow or database annotation (referred here simply as Reference_Files) used in this project. This document describes the inputs/outputs needed and where to locate the information distributed among different person or different directories on a digital infrastructure as presented in figure 3 but with more details to improve the reproducibility of the DRO ^33^.

The markdown format was kept as it was easy to write/read by a human or to generate via a python script. This file was formatted into a simplified RDF-like triples structure, were each files-objects (subject) were linked to information (objects) by a property. This format allowed to use the following properties to describe each file-objects we had: The file-object name or identification, a brief description (i.e. about the software used or the data content), the file-object version, the input(s)/output(s), the associated documents, hyperlink(s) to remote database or citation, the location of the file-object on the project directory or archiving system, and the author(s) to contact for questions. These associations could be viewed as a graph to display the important files and pipelines used. This document was useful to understand how exactly the different files were generated, and to recover the scripts and input/ouput that were used, even after prolonged periods and to use them again, which permit for example to reproduce data with novel or updated annotation files. Furthermore, if an error was detected within a script, the results and figures downstream that needed to be recomputed could be easily found. This documentation file was made available on gitlab (Data Citation 4, *gitlab* Systems_Genetics_of_Sleep_Regulation).

### Data Mining Website

The DRO built for this systems genetics resource is constituted of the following collection: raw-data, processed data, Rmarkdown reports, results & interpretation, workflow, scripts, and metadata. To improve the reproducibility of our integrative visualization method (see HivePlots below), we provided some data-mining tools, a server to store some intermediate results, and a web application ^34^ (Data Citation 3, bxd.vital-it.ch https://bxd.vital-it.ch/#/dataset/). The home page of the web application displays the information for the NREM sleep gain during the 24 hours (in four 6-hour intervals) after sleep deprivation. Three data-mining tutorials were described on the website the web interface to: (i) mine a single phenotype, (ii) search for a gene, and (iii) compare hiveplots. Currently, no centralized repository exists containing all types of phenotypic data that were extracted within this project. This web-interface can, however be viewed as a hub for this DRO that became findable and accessible with a web-browser. With this web resource, we provided an advanced interactive interface for EEG/behavioral end-phenotypes and their associated intermediate phenotypes (variants, metabolites, gene expression). Compared to other web-resources for systems genetics like GeneNetwork where the principal focus is QTL mining, this interface provides an integrative view of this one dataset, with also data files and link to code to reproduce some of our analyses in the form of Rmarkdown, like the prioritization strategy.

### Low-Layer Analyses

#### Sleep-wake state annotation

To assist the annotation of this extensive dataset (around 20 million 4 s epochs), we developed a semiautomated scoring system. The 4-day recordings of 43 mice (19% of all recordings), representing animals from 12 strains, were fully annotated visually by an expert according to established criteria ^22^. Due to large between-line variability in EEG signals, even after normalization, a partial overlap of the different sleep-wake states remained, as evidenced by the absolute position of the center of each state cluster, which differed even among individuals of the same line (precluding the use of 1 “reference” mouse), even per line, to reliably annotate sleep-wake states for the others. To overcome this problem, 1 day out of 4 (i.e., Day 3 or R1, which includes the SD) was visually annotated for each mouse. These 4 seconds sleep-wake scores were used to train the semiautomatic scoring algorithm, which took as input 82 numerical variables derived from the analyses of EEG and EMG signals using frequency-(discrete Fourier transform [DFT]) and time-domain analyses performed at 1 second resolution. We then used these data to train a series of support vector machines (SVMs)^35^ specifically tailored for each mouse, using combinations of the 5 or 6 most informative variables out of the 82 input variables. The best-performing SVMs for a given mouse were then selected based on the upper-quartile performance for global classification accuracy and sensitivity for REM sleep (the sleep-wake state with the lowest prevalence) and used to predict sleep-wake states in the remaining 3 days of the recording. The predictions for 4 consecutive 1-s epochs were converted into 1 four-second epoch. Next, the results of the distinct SVMs were collapsed into a consensus prediction, using a majority vote. In case of ties, epochs were annotated according to the consensus prediction of their neighboring epochs. To prevent overfitting and assess the expected performance of the predictor, only 50% of the R1 manually annotated data from each mouse were used for training. The classification performance was assessed by comparing the automatic and visual scoring of the fully manually annotated 4 d recordings of 43 mice. The global accuracy was computed using a confusion matrix ^36^ of the completely predicted days (B1, B2, and R2). For all subsequent analyses, the visually annotated Day 3 (R1) recording and the algorithmically annotated days (B1, B2, and R2) were used for all mice, including those for which these days were visually annotated. The resulting sleep-wake state annotation together with EEG power spectra and EMG levels were saved as binary files (.smo) with their corresponding metadata files (.hdr) and deposited at FigShare (Data Citation 1: Figshare https://…). For more information on *.smo* and *.hdr* files, see Usage Notes.

#### EEG/Behavioral Phenotyping

We quantified 341 phenotypes based on the sleep-wake states, LMA, and the spectral composition of the EEG, constituting 3 broad phenotypic categories. For the first phenotypic catergory (“State”), the 96 hours sleep-wake sequence of each animal was used to directly assess traits in 3 “state”-related phenotypic subcategories: (i) duration (e.g., time spent in wakefulness, NREM sleep, and REM sleep, both absolute and relative to each other, such as the ratio of time spent in REM versus NREM sleep); (ii) aspects of their distribution over the 24 h cycle (e.g., time course of hourly values, midpoint of the 12 h interval with highest time spent awake, and differences between the light and dark periods); and (iii) sleep-wake architecture (e.g., number and duration of sleep-wake bouts, sleep fragmentation, and sleep-wake state transition probabilities). Similarly, for the second phenotypic category (“LMA”) overall activity counts per day, as well as per unit of time spent awake, and the distribution of activity over the 24 h cycle was extracted from the LMA data. As final phenotypic category (“EEG”), EEG signals of the 4 different sleep-wake states (wakefulness, NREM sleep, REM sleep, and theta-dominated waking [TDW], see below) were quantified within the 4-s epochs matching the sleep-wake states using DFT (0.25 Hz resolution, range 0.75–90 Hz, window function Hamming). Signal power was calculated in discrete EEG frequency bands—i.e., delta (1.0–4.25 Hz, δ), slow delta (1.0–2.25 Hz; δ1), fast delta (2.5–4.25; δ2), theta (5.0–9.0 Hz; θ), sigma (11–16 Hz; σ), beta (18–30 Hz; β), slow gamma (32–55 Hz; γ1), and fast gamma (55–80 Hz; γ2). Power in each frequency band was referenced to total EEG power over all frequencies (0.75–90 Hz) and all sleep-wake states in days B1 and B2 to account for interindividual variability in absolute power. The contribution of each sleep-wake state to this reference was weighted such that, e.g., animals spending more time in NREM sleep (during which total EEG power is higher) do not have a higher reference as a result ^37^. Moreover, the frequency of dominant EEG rhythms was extracted as phenotypes, specifically that of the theta rhythm characteristic of REM sleep and TDW. The latter state, a substate of wakefulness, defined by the prevalence of theta activity (6.0–10.0 Hz) in the EEG during waking ^38,39^, was quantified according to the algorithm described in ^40^. We assessed the time spent in this state, the fraction of total wakefulness it represents, and its distribution over 24 h. Finally, discrete, paroxysmal events were counted, such as sporadic spontaneous seizures and neocortical spindling, which are known features of D2 mice ^41^, which we also found in some BXD lines.

All phenotypes were quantified in baseline and recovery separately, and the effect of SD on all variables was computed as recovery versus baseline differences or ratios. The recovery-to-baseline contrasts are the focus of this paper. Obviously, some of the 341 phenotypes are strongly correlated (e.g., the time spent awake and asleep in a given recording interval), resulting in identical QTLs (albeit with different association strengths). Pascal source code used for EEG/behavioral phenotyping was made available on gitlab (Data Citation 4, *gitlab* Systems_Genetics_of_Sleep_Regulation). Processed phenotypes and descriptions were made available at bxd.vital-it.ch (Data Citation 3, bxd.vital-it.ch https://bxd.vital-it.ch/#/dataset/).

#### Read alignment

For gene expression quantification, we used a standard pipeline that was already applied in a previous study ^6^. Bad quality reads tagged by Casava 1.82 were filtered from fastq files and reads were mapped to MGSCv37/mm9 using the STAR splice aligner (v 2.4.0g) with the *2pass* pipeline ^42^.

#### Genotyping

The RNA-seq dataset was also used to complement the publicly available GeneNetwork genetic map (www.genenetwork.org), thus increasing its resolution. RNA-seq variant calling was performed using the Genome Analysis ToolKit (GATK) from the Broad Institute, using the recommended workflow for RNA-seq data ^43^. To improve coverage depth, 2 additional RNA-seq datasets from other projects using the same BXD lines were added ^6^. In total, 6 BXD datasets from 4 different tissues (cortex, hypothalamus, brainstem, and liver) were used. A hard filtering procedure was applied as suggested by the GATK pipeline ^43–45^. Furthermore, genotypes with more than 10% missing information, low quality (<5,000), and redundant information were removed. GeneNetwork genotypes, which were discrepant with our RNA-seq experiment, were tagged as “unknown” (mean of 1% of the GeneNetwork genotypes/strain [0.05% ≤n≤ 8%]). Finally, GeneNetwork and our RNA-seq genotypes were merged into a unique set of around 11,000 genotypes, which was used for all subsequent analyses. This set of genotypes was already used successfully in a previous study of BXD lines ^6^ and is available through our “Swiss-BXD” web interface (Data Citation 3, bxd.vital-it.ch https://bxd.vital-it.ch/#/dataset/).

#### Protein damage prediction

Variants detected by our RNA-seq variant calling were annotated using Annovar ^46^ with the RefSeq annotation dataset. Nonsynonymous variations were further investigated for protein disruption using Polyphen-2 version 2.2.2 ^47^, which was adapted for use in the mouse according to recommended configuration. Variant annotation file and polyphen2 scores were made available here (Data Citation 5, *figshare* https://figshare.com/s/51916157a22357755de8).

#### Gene expression quantification

Count data was generated using htseq-count from the HTseq package using parameters “stranded = reverse” and “mode = union” ^48^. Gene boundaries were extracted from the mm9/refseq/reflat dataset of the UCSC table browser. Raw counts were made available here (Data Citation 5, *figshare* https://figshare.com/s/51916157a22357755de8).

### Intermediate-Layer Analyses

#### Gene expression normalization

EdgeR was then used to normalize read counts by library size. Genes with with low expression value were excluded from the analysis, and the raw read counts were normalized using the TMM normalization ^49^ and converted to log counts per million (CPM). R. Although for both tissues, the RNA-seq samples passed all quality thresholds, and among-strain variability was small, more reads were mapped in cortex than in liver, and we observed a somewhat higher coefficient of variation in the raw gene read count in liver than in cortex. Genes expression as CPM or log2 CPM were made available here: (Data Citation 5, *figshare* https://figshare.com/s/51916157a22357755de8).

#### Differential expression

To assess the gene differential expression between the sleep-deprived and control conditions, we used the R package limma ^50^ with the voom weighting function followed by the limma empirical Bayes method ^51^. Differential expression tables were made available here: (Data Citation 5, *figshare* https://figshare.com/s/51916157a22357755de8).

#### QTL mapping

The R package qtl/r ^28^ was used for interval mapping of behavioral/EEG phenotypes (phQTLs) and metabolites (mQTLs). Pseudomarkers were imputed every cM, and genome-wide associations were calculated using the Expected-Maximization (EM) algorithm. p-values were corrected for FDR using permutation tests with 1,000 random shuffles. The significance threshold was set to 0.05 FDR, a suggestive threshold to 0.63 FDR, and a highly suggestive threshold to 0.10 FDR according to ^52,53^. QTL boundaries were determined using a 1.5 LOD support interval. To preserve sensitivity in QTL detection, we did not apply further p-value correction for the many phenotypes tested. Effect size of single QTLs was estimated using 2 methods. Method 1 does not consider eventual other QTLs present and computes effect size according to 1 − 10^(−(2/n)*LOD). Method 2 does consider multi-QTL effects and computes effect size by each contributing QTL by calculating first the full, additive model for all QTLs identified and, subsequently, estimating the effects of each contributing QTL by computing the variance lost when removing that QTL from the full model (“drop-one-term” analysis). For Method 2, the additive effect of multiple suggestive, highly suggestive, and significant QTLs was calculated using the fitqtl function of the qtl/r package ^54^. With this method, the sum of single QTL effect estimation can be lower than the full model because of association between genotypes. In the Results section, Method 1 was used to estimate effect size, unless specified otherwise. It is important to note that the effect size estimated for a QTL represents the variance explained of the genetic portion of the variance (between-strain variability) quantified as heritability and not of the total variance observed for a given phenotype (i.e., within-plus between-strain variability).

For detection of eQTLs, cis-eQTLs were mapped using FastQTL ^28^ within a 2 Mb window for which adjusted p-values were computed with 1,000 permutations and beta distribution fitting. The R package qvalue ^55^ was then used for multiple-testing correction as proposed by ^28^. Only the q-values are reported for each cis-eQTL in the text. Trans-eQTL detection was performed using a modified version of FastEpistasis ^29^, on several million associations (approximately 15,000 genes × 11,000 markers), applying a global, hard p-value threshold of 1E−4.

List of ph-QTLs, cis-eQTL, trans-eQTL and m-QTLs were made available here: (Data Citation 5, *figshare* https://figshare.com/s/51916157a22357755de8).

### High-layer Analyses

#### Hiveplot visualization

Hiveplots were constructed with the R package HiveR^56^ for each phenotype. Gene expression and metabolite levels represented in the hiveplots come from either the “Ctr” (control) or SD molecular datasets according to the phenotype represented in the hiveplot; i.e., the “Ctr” dataset is represented for phenotypes related to the baseline (“bsl”) condition, while the SD dataset is shown for phenotypes related to recovery (“rec” and “rec/bsl”). For a given hiveplot, only those genes and metabolites were included (depicted as nodes on the axes) for which the Pearson correlation coefficient between the phenotype concerned and the molecule passed a data-driven threshold set to the top 0.5% of all absolute correlations between all phenotypes on the one hand and all molecular (gene expression and metabolites) on the other. This threshold was calculated separately for “Bsl” phenotypes and for “Rec” and “Rec/Bsl” phenotypes and amounted to absolute correlation thresholds of 0.510 and 0.485, respectively. The latter was used for the recovery phenotypes. Associations between gene expressions and metabolites represented by the edges in the hiveplot were filtered using quantile thresholds (top 0.05% gene–gene associations, top 0.5% gene–metabolite associations). We corrected for cis-eQTL confounding effects by computing partial correlations between all possible pairs of genes. Hiveplots figures and Rmarkdowns reports were made available here (Data Citation 5, *figshare* https://figshare.com/s/51916157a22357755de8).

#### Candidate-gene prioritization strategy

In order to prioritize genes in identified QTL regions, we chose to combine the results of the following analyses: (i) QTL mapping (phQTL or mQTL), (ii) correlation analysis, (iii) expression QTL (eQTL), (iv) protein damaging–variation prediction, and (v) DE. Each result was transformed into an “analysis score” using a min/max normalization, in which the contribution of extreme values was reduced by a winsorization of the results. These analysis scores were first associated with each gene (see below) and then integrated into a single “integrated score” computed separately for each tissue, yielding 1 integrated score in cortex and 1 in liver. The correlation analysis score, eQTL score, DE score, and protein damaging– variation score are already associated to genes, and these values were therefore attributed to the corresponding gene. To associate a gene with the ph-/m-QTL analysis score (which is associated to markers), we used the central position of the gene to infer the associated ph-/m-QTL analysis score at that position. In case of a cis-eQTL linked to a gene or a damaging variation within the gene, we used the position of the associated marker instead. To emphasize diversity and reduce analysis score information redundancy, we weighted each analysis score using the Henikoff algorithm. The individual scores were discretized before using the Henikoff algorithm, which was applied on all the genes within the ph-/mQTL region associated with each phenotype. The integrated score was calculated separately for cortex and liver. We performed a 10,000-permutation procedure to compute an FDR for the integrated scores. For each permutation procedure, all 5 analysis scores were permutated, and a novel integrated score was computed again. The maximal integrated score for each permutation procedure was kept, and a significance threshold was set at quantile 95. Applying the Henikoff weighting improved the sensitivity of the gene prioritization. E.g., among the 91 behavioral/EEG phenotypes quantified with 1 or more suggestive/significant QTL after SD, 40 had at least 1 gene significantly prioritized with Henikoff weighting, against 32 without. Gene prioritization figures and Rmarkdown reports were made available here (Data Citation 5, *figshare* https://figshare.com/s/51916157a22357755de8)

### Reproducibility of the pipeline

#### Technical reproducibility of the pipeline

To assess the reproducibility of our analytical pipeline, we asked a bioinformatician that was not involved in the data collection and analysis to reanalyze some of the results. A relatively short computational time as well as importance in the published results were taken as selection criteria of analyses to be replicated. The TMM normalisation of RNA-seq counts, differential gene expression, cis-eQTL detection, and the ph-/m-QTL mapping for 4 sleep phenotypes (slow delta power gain after SD, fast delta power after SD, theta peak frequency shift after SD and NREM sleep gain in the dark after SD) and 2 metabolites (Phosphatidylcholine ae C38:2 and alpha amino-adipic acid) used as main examples in our previous publication were all re-analyzed. Finally, gene prioritization and hiveplot visualization of these 4 examples were replicated. Originally, ties in the nodes ranking function on the hiveplots axis was solved using the “random” method, but this function was modified in the hiveplot code and set as “first” to remain deterministic (see Technical Validation for results).

#### Reanalysis with mm10

To quantify the effect of new standards and robustness of our end-results and interpretation we changed some analyses within our low layer. The mm10 genome assembly was set as our new reference and the gene expression was reanalysed from the raw fastq files with the BioJupies reproducible pipeline ^57^ ^58^ that use kallisto pseudo-alignement ^59^. The gene positions were retrieved from the headers of the ENSEMBL fasta file used by BioJupies (Mus_musculus.GRCm38.cdna.all.fa.gz). Genotypes were downloaded from GeneNetwork database and our annovar/polyphen2 variations positions based on mm9 were adapted to mm10 using CrossMap version 0.2.4 ^60^. The analyses performed to assess the technical reproducibility of our pipeline (see above) were finally replicated using these new files. (see Technical Validation for results).

## Data Records

EEG/EMG power spectra and locomotor activity files were submitted to *figshare* Data Citation 1: *Figshare* LinkWithSubmission). Raw data of RNA-sequencing were submitted to Gene Expression Omnibus (Data Citation 2: *NCBI Gene Expression Omnibus* GSE112352). Processed phenotypes files as gene expression, metabolites level and mean EEG/behavioral phenotypes per lines, as well as phenotypes descriptions, were submitted to our data-mining web-site (Data Citation 3, bxd.vital-it.ch https://bxd.vital-it.ch/#/) on the ‘Downloads’ panel. Scripts and code were submitted to gitlab Data Citation 4, *gitlab* Systems_Genetics_of_Sleep_Regulation). Intermediate files required to run these scripts were submitted to *figshare* (Data Citation 5, *figshare* https://figshare.com/s/51916157a22357755de8). BXD genotypes are available on GeneNetwork (Data Citation 6, *GeneNetwork*, GN600).

## Technical Validation

### Compare genotype RNA-seq vs GeneNetwork

To verify the genetic background of each mice we phenotyped, we analyzed the correspondence between GeneNetwork genotypes and RNA-seq variants detected by GATK. For the 3811 GeneNetwork genotypes, 1289 could be recalled in our RNA-seq variant calling pipeline. Figure 4 shows the similarity proportion between RNA-seq variants and GeneNetwork genotypes, for each pair of BXD lines. Our BXD63 was more similar with the GeneNetwork BXD67 than with the BXD63, probably due to mislabeling. We therefore chose to exclude this line. The matrix also shows the genetic similarity between BXD73 and BXD103 (now renamed as BXD73b), between BXD48 and BXD96 (now BXD48a) and between BXD65 and BXD97 (now BXD65a), which confirmed the renaming of these BXD lines on GeneNetwork.

**Figure 4:**
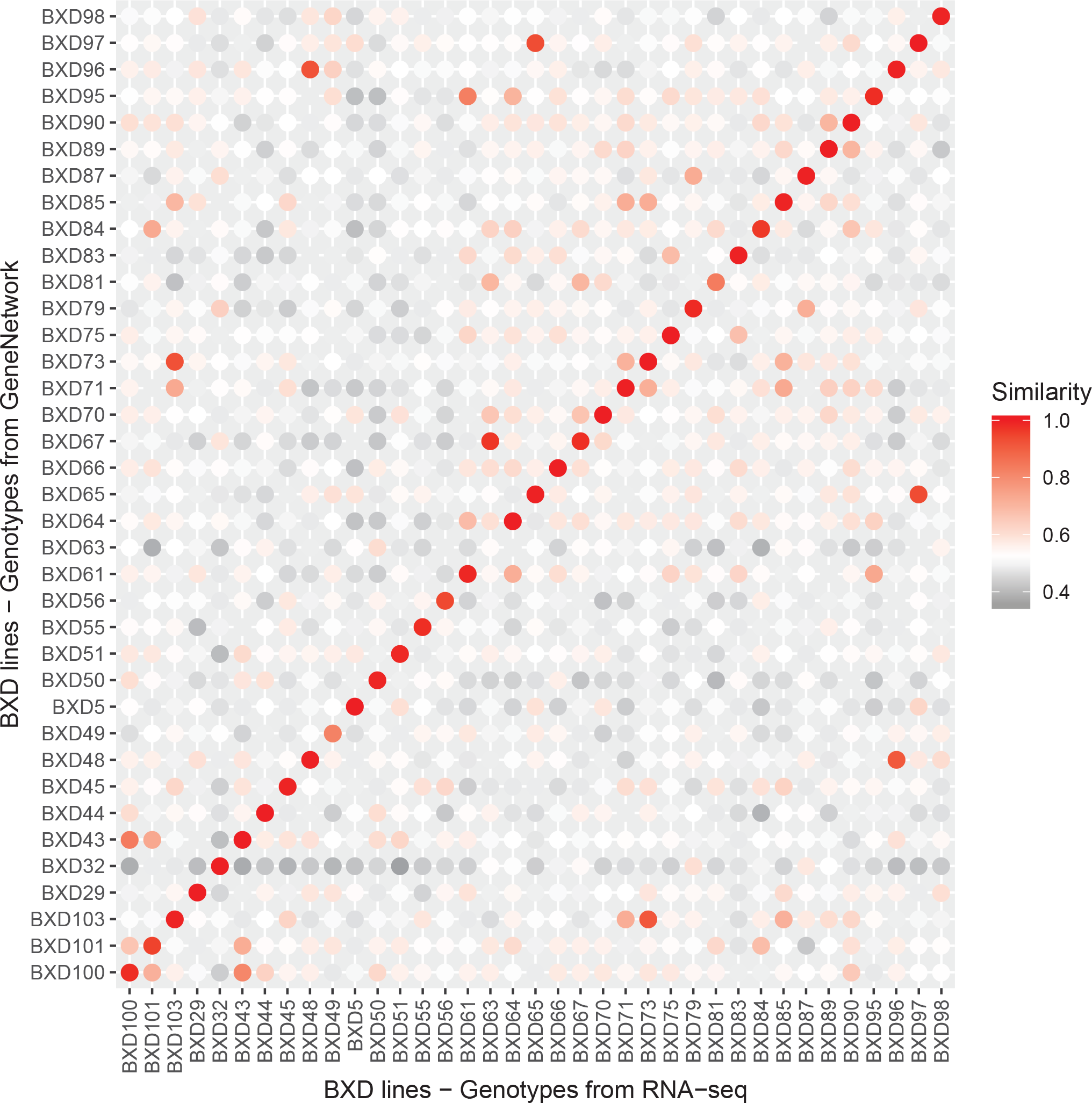
Similarity matrix [in %] between RNA-seq variant calling and GeneNetwork genotypes. A similarity of 1 indicates that all common genotypes are similar. We here compare only genotypes that were labeled as ‘B’ or ‘D’ and excluded unknown ‘U’ or heterozygous ‘H’ genotypes.

### Reproducibility of the pipeline

#### Technical reproducibility of the pipeline

To assess the technical reproducibility of the pipeline, a bioinformatics student (NG) new to the project reproduced chosen steps of the bioinformatic pipeline. The results (Figure 5, upper part) were consistent with previous analyses (PLOS paper figures: 2C, 4C left, 7D, and 7C bottom). The robustness of the pipeline was verified because the same conclusions could be drawn. For examples, the same 3 genes showed the largest differential expression after SD in the cortex (*Arc*, *Plin4*, and *Egr2* in Figure 5B). Moreover, the *Acot11* gene was prioritized by gene prioritization (Figure 5D&E). Nevertheless, the numbers of significant genes of cis eQTL showed variations compared to previous analysis {Diessler, 2018 #495} due to use of significance threshold for visualization. For example, the number of genes with significant QTL unique to Cortex SD changed from 870 (PLOS paper Figure 2C) to 872 (Figure 5A). The genes are considered as significant if their FDR-adjusted p-value is below or equal to 0.05, which is obtained by estimating the β-distribution fitting of random permutations tests. Changing the fastqtl version (version 1.165 to version 2.184) seems to change the pseudo-random number generation, even when using the concept of fixed seed. Consequently, the number of genes considered as significant varies because their FDR-adjusted p-value passes just above or below the threshold (FDR in the range of 0.04864 to 0.05054). This confirms that looking at the order of magnitude is important, though the use of significance threshold is convenient.

**Figure 5:**
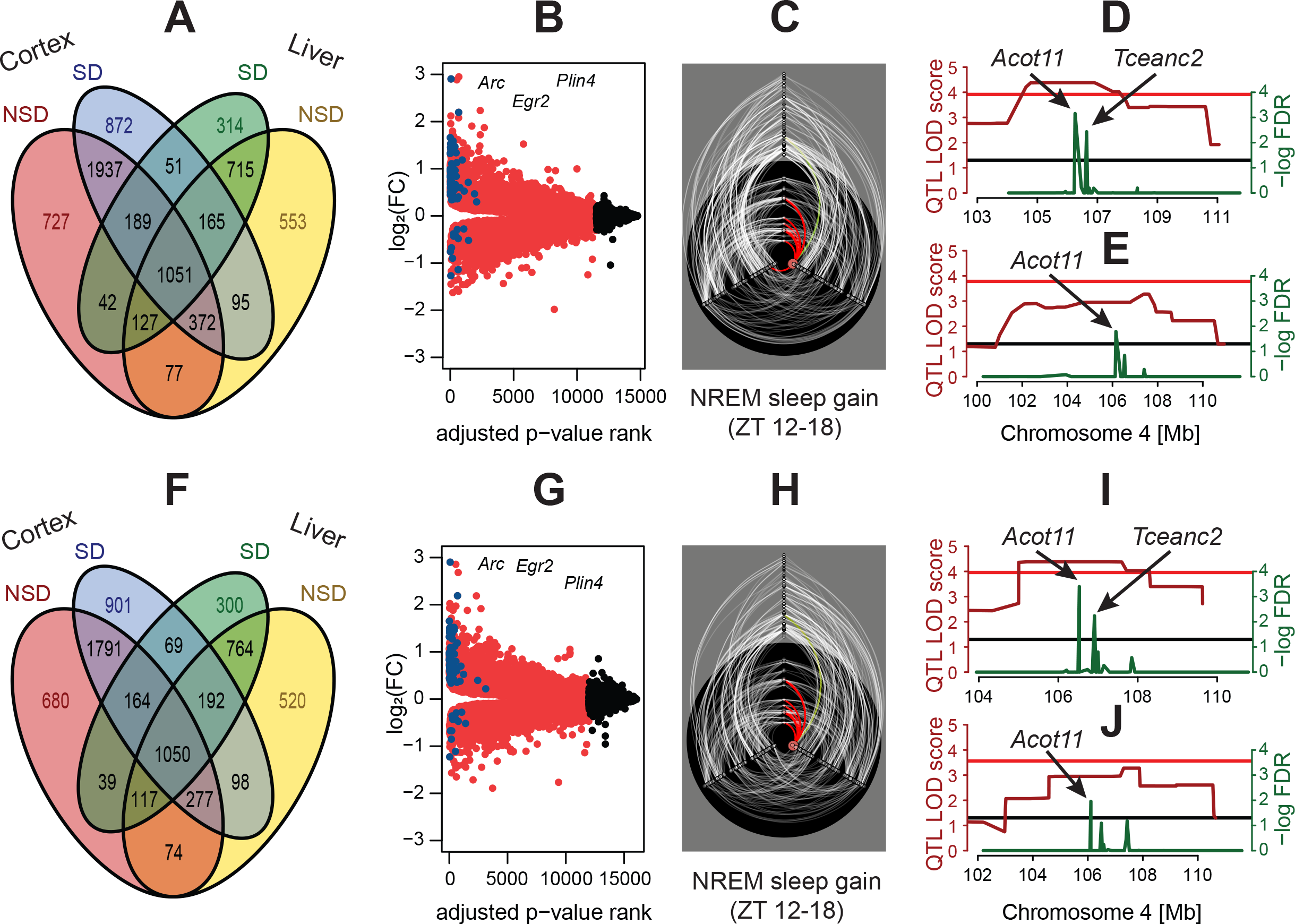
Robustness of the analysis pipeline. A to E: Technical reanalysis with mm9 reference genome. F to J: Reanalysis with mm10 reference genome. A and F: Venn diagram of significant cis-eQTL. B and G: Volcano plot of differential gene expression in cortex. C and H: Hiveplot NREM sleep gain during recovery of with highlight on Acot11. D, E, I, and J: Gene prioritization for NREM sleep gain during recovery (D and I) or phosphatidylcholine acyl-alkyl C38:2 levels (E and J). recovery=first 6 hours of dark period after sleep deprivation (ZT 12-18), SD=sleep deprivation, NSD=not sleep deprivation (control), FC= fold-change, NREM=non-REM, REM=Rapid eye movement, LOD=logarithm of odds, FDR=false discovery rate.

Moreover, the reanalysis process helped to improve the code documentation by explicitly writing project-related knowledge, such as common abbreviations. Having another perspective on the code also allowed to improve its structure. Indeed, a retrospective overview helped improve the organisation of files, which was more difficult to do within the implementation of the project because the code was incrementally created and adapted. The process allowed to catch and correct minor mistakes or make improvements improved readability and consistency. For example, it was highlighted that the ranking function used in hiveplot to order nodes in the axes was using the “random” argument for differentiating ties. As a key concept of the hiveplots was to be fully reproducible in the sense of “perpetual uniformity” ^56^, we changed the ties.method parameter to “first” so that the same input always gives the same result, without having to fix a seed for the pseudo-random generation. Another example was the ranking of the x-axis in the gene DE volcano plot and the colouring that were based on log-odds values (B statistic according to in limma R package) instead of FDR-adjusted p-values. However, this reproducibility ‘experiment’ was internal to the group, which facilitate communication such as which steps to focus on and whether to run them locally or in a high-performance computing (HPC) structure. An assessment of the computational requirements for each step, such as computing time, memory, software, and libraries used may be interesting to provide to facilitate external reproducibility.

#### Reanalysis with mm10

To assess the influence of the reference genome used in the analyses, we reproduced selected parts of bioinformatic pipeline using the updated version (mm10 instead of mm9). The results (Figure 5, lower part, table 2 and 3) were consistent with previous analyses but presented also some substantial variations. The cis-eQTL detection revealed differences in the number of significant associations found, as showed in Table 2. These differences could be mainly explained by small q-value variation around the significant threshold. Nevertheless, around 5% of cis-eQTLs did not reproduce even at a more permissive significant threshold (0.1 FDR), which affected some of our end results. For example, *Wrn* was no longer prioritized for the gain of slow EEG delta power (δ1) after SD compared to previous results on mm9. Although the cis-eQTL for *Wrn* was present in both assemblies for the ‘Cortex Control’ samples, it disappeared for ‘Cortex SD’ samples using mm10. A number of factors could have contributed to this discrepancy among which i) the variations between mm9 and mm10 could change the mappability of some transcripts, although this did not seem to be the case for the *Wrn* sequence, ii) pseudo-alignment (Kallisto) was used instead of alignment (STAR), which may have influenced the quantification, iii) bad quality reads were filtered with our STAR pipeline according to Casava 1.82 but not with Kallisto, and iv) variant calling on RNA-seq data to add markers was not done for mm10, so only markers from GeneNetwork were used. Specifically to the latter factor, the marker closest to *Wrn* gene in mm9 merged genotypes (rs51740715) is not present in mm10. The change in the number of genetic markers could have therefore influenced the cis-eQTL detection, which is an important in the gene prioritization that resulted in the identification of *Wrn* as candidate underlying the EEG delta power (δ1) trait under mm9.

**Table 2:** Comparison of cis-eQTL summary statistics in mm9 and mm10 reanalyses. ‘Unique’ is defined as specific to an assembly (mm9 or mm10). Significance is defined as a q-value below or equal to 0.05. ‘Overlapping’ is defined as common between mm9 and mm10 reanalyses. ‘Almost overlapping’ is defined as uncommon between mm9 and mm10 that would be common if a threshold of 0.1 was used instead of 0.05.

|  | Liver NSD |  | Liver SD |  | Cortex NSD |  | Cortex SD |  |
| --- | --- | --- | --- | --- | --- | --- | --- | --- |
| Assembly | mm9 | mm10 | mm9 | mm10 | mm9 | mm10 | mm9 | mm10 |
| Total genes | 14103 | 12647 | 14103 | 12647 | 14889 | 15734 | 14889 | 15734 |
| Unique genes | 2405 | 949 | 2405 | 949 | 1043 | 1888 | 1043 | 1888 |
| Genes with significant cis-eQTL | 3155 | 3092 | 2654 | 2695 | 4522 | 4192 | 4732 | 4542 |
| Proportion of genes with significant cis-eQTL | 0.22 | 0.24 | 0.19 | 0.21 | 0.30 | 0.27 | 0.32 | 0.29 |
| Genes with significant cis-eQTL overlapping | 2255 |  | 1911 |  | 3204 |  | 3483 |  |
| Genes with not significant cis-eQTL overlapping | 8375 |  | 8857 |  | 9062 |  | 8801 |  |
| Genes with significant cis-eQTL not overlapping | 900 | 837 | 743 | 784 | 1318 | 988 | 1249 | 1059 |
| Genes with significant cis-eQTL almost overlapping | 2995 | 2898 | 2535 | 2505 | 4201 | 4019 | 4441 | 4350 |

**Table 3:** Comparison of gene DE in mm9 and mm10 reanalyses. Suggestive is defined as a q-value below or equal to 0.1.

|  | Liver |  | Cortex |  |
| --- | --- | --- | --- | --- |
|  | mm9 | mm10 | mm9 | mm10 |
| Total genes | 12539 | 13264 | 14754 | 16057 |
| DE genes | 7573 | 8754 | 11534 | 11980 |
| Proportion of DE genes | 0.6040 | 0.6600 | 0.7818 | 0.7461 |
| Suggestive DE genes | 8253 | 9392 | 12069 | 12580 |
| Proportion of suggestive DE genes | 0.6582 | 0.7081 | 0.8180 | 0.7835 |

## Usage Notes

### SMO files

Binary *.smo* files were structured as follows: Each file contains a 4-day recording or precisely 86’400 consecutive 4s epochs. Each 4s epoch contains the following information: one byte character and 404 single precision floating-points, which represent, respectively: sleep-wake state of the 4s epoch as a character (wake = ‘w’, NREM sleep = ‘n’, REM sleep = ‘r’, wake w/ EEG artifact = ‘1’, NREM sleep w/ EEG artifact = ‘2’, REM sleep w/ EEG artifact = ‘3’, wake w/ spindle-like EEG activity = ‘4’, NREM sleep w/ spindle-like EEG activity = ‘5’, REM sleep w/ spindle-like EEG activity = ‘6’, Paroxysmal EEG activity in wake = ‘7’, Paroxysmal EEG activity in NREM sleep = ‘8’, Paroxysmal EEG activity in REM sleep = ‘9’), EEG power density from the full DFT spectrum of the 4s epoch from 0.00 Hz to 100.00 Hz (401 values at 0.25-Hz resolution), the EEG variance, the EMG variance, and temperature. Temperature was not measured and was set to 0.0.

### HDR files

Text *.hdr* files are generated alongside their corresponding *.smo* file and contain among other information, the mouse ID (*Patient*) and recording date.

### Rmarkdown scripts

Some of the Rmarkdown scripts were created for a remote cluster environment on a CentOS distribution which required the use of a second script that generated the document with the *rmarkdown::render()* function and pass the expected function arguments. Therefore some functions that use the parallel package in R are only executable on a linux environment (i.e. mclapply()). These functions can be modified with the *doSNOW* R library to be applicable on a windows environment. The author can set many option in the YAML (Yet Another Markup Language) header to: create dynamic and readable table that contains multiple rows, hide/show source code or integrated CSS style and table of contents. The reports can be visualized using any web-browser.

## Acknowledgements

We thank Yann Emmenegger, Mathieu Piguet and Josselin Soyer for the help organizing and handling the BXD mice; Benita Middleton and Debra J. Skene at the university of Surrey for metabolomics quantification; the experts at the Lausanne Genomics Technologies Facility (GTF) for their help with RNA-sequencing; Nicolas Guex, Mark Ibberson, Frederic Burdet, Alan Bridge, Lou Götz, Marco Pagni, Martial Sankar and Robin Liechti for their bioinformatics expertise and help.

Computations and analyses were performed at Vital-IT (http://www.vital-it.ch).

PF, SD, and MJ were funded by the University of Lausanne (Etat de Vaud). MJ and NG were funded by the Swiss National Science Foundation grant to PF (31003A_173182). MJ and SD were funded by the Swiss National Science Foundation grant to PF and IX (CRSII3_13620).

## Author contributions

M.J. Data analysis, data organization, manuscript writing

N.G. Pipeline reproduction, data organization, manuscript writing

S.D. Data generation and supervision for experiment 1 and 2

P.F. Data analysis, design experiments 1 and 2, project supervision, manuscript writing

I.X. Conceptualization, methodology, project supervision, manuscript writing

